# Discovery of a 382-nt deletion during the early evolution of SARS-CoV-2

**DOI:** 10.1101/2020.03.11.987222

**Authors:** Yvonne CF Su, Danielle E Anderson, Barnaby E Young, Feng Zhu, Martin Linster, Shirin Kalimuddin, Jenny GH Low, Zhuang Yan, Jayanthi Jayakumar, Louisa Sun, Gabriel Z Yan, Ian H Mendenhall, Yee-Sin Leo, David Chien Lye, Lin-Fa Wang, Gavin JD Smith

## Abstract

To date, the SARS-CoV-2 genome has been considered genetically more stable than SARS-CoV or MERS-CoV. Here we report a 382-nt deletion covering almost the entire open reading frame 8 (ORF8) of SARS-CoV-2 obtained from eight hospitalized patients in Singapore. The deletion also removes the ORF8 transcription-regulatory sequence (TRS), which in turn enhances the downstream transcription of the N gene. We also found that viruses with the deletion have been circulating for at least four weeks. During the SARS-CoV outbreak in 2003, a number of genetic variants were observed in the human population [1], and similar variation has since been observed across SARS-related CoVs in humans and bats. Overwhelmingly these viruses had mutations or deletions in ORF8, that have been associated with reduced replicative fitness of the virus [2]. This is also consistent with the observation that towards the end of the outbreak sequences obtained from human SARS cases possessed an ORF8 deletion that may be associated with host adaptation [1]. We therefore hypothesise that the major deletion revealed in this study may lead to an attenuated phenotype of SARS-CoV-2.

---

On 1 December 2019, a novel coronavirus emerged from Hubei province in China and infected people visiting the Huanan seafood market in Wuhan [3]. The virus demonstrated efficient human-to-human transmission within mainland China and subsequently spread across many countries. The virus was soon identified as 2019-nCoV, more recently designated SARS-CoV-2, while the disease is referred to as COVID-19. On 30 January 2020, the World Health Organization declared a Public Health Emergency of International Concern. As of 10 March 2020, the COVID-19 outbreak has led to 113,702 confirmed cases and 4,012 deaths globally [4]. Zoonotic viruses have crossed the species barrier from animals to humans and the success of an interspecies transmission is the ability of a novel virus to adapt, via genetic mutation events, to a new host and cause sustained transmission that can lead to a significant outbreak.

Nasopharyngeal swabs collected from hospitalized patients positive for SARS-CoV-2 in Singapore were subjected to next generation sequencing (NGS) analysis, with and without passaging in Vero-E6 cells. The NGS data revealed a 382-nt deletion towards the 3’ end of the viral genomes obtained from multiple patients (Suppl. Table 1). To confirm this observation, specific PCR primers were designed flanking the deleted region and Sanger sequencing performed (Suppl. Fig. 1). This verified the deletion at positions 27,848 to 28,229 of the SARS-CoV-2 genome. Interrogation of the NGS assemblies of these 382-nt deletion variants (referred to hereafter as Δ382) indicated that the virus populations were homogenous.

Apart from the 382-nt deletion, the genome organization of the Δ382 viruses is identical to that of other SARS-CoV-2 (Fig. 1A). Closer examination of the deletion indicated that it spans an area of the ORF7b and ORF8 that includes the ORF8 transcriptional regulator sequence (TRS), eliminating ORF8 transcription (Fig. 1B). The ORF8 region has been identified as an evolutionary hotspot of SARSr-CoVs. For instance, sequences of human SARS-CoV TOR2 and LC2 exhibit 29-nt and 415-nt deletions, whereas those of bat SARS-CoV JTMC15 have discontinuous deletions in ORF7/8 (Fig. 1B) [5-8]. The ORF8 region of SARS-CoVs has been shown to play a significant role in adaptation to the human host following interspecies transmission [7] and virus replicative efficiency [2]. Similarly the disruption of ORF8 region in Δ382 viruses may be a result of human adaptation after the emergence of SARS-CoV-2.

**Figure 1.**
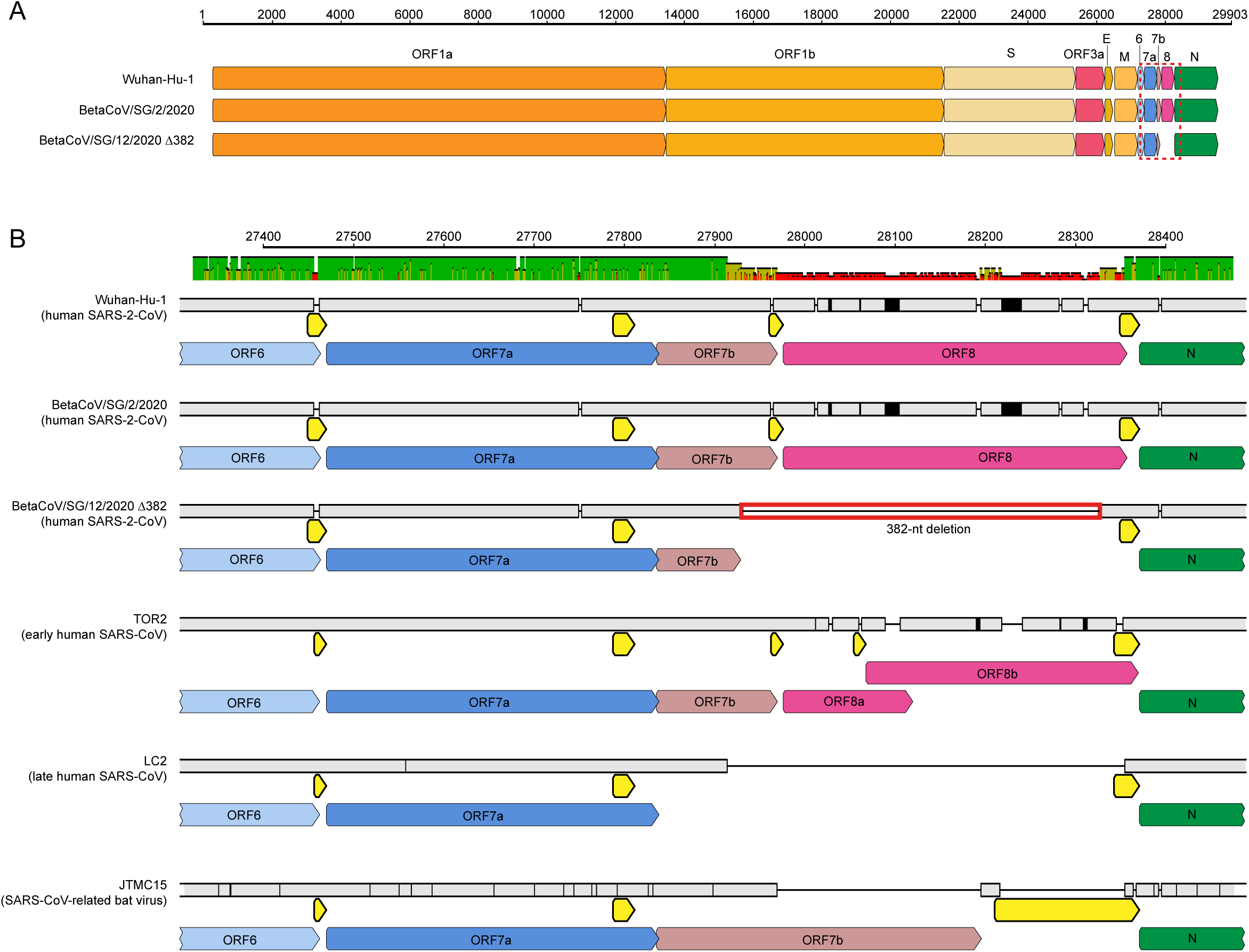
Schematic comparison of SARS-2-CoV, SARS-CoV and SARSr-CoV genomes. **A.** Full genome sequences of SARS-2-CoV isolates Wuhan-Hu-1 (GenBank: MN908947), BetaCoV/Singapore/2/2020 (GISAID: 407987), and BetaCoV/Singapore/12/2020 Δ382 (GISAID: EPI_ISL_414378). **B.** Magnification of genomic region (dashed box in Figure 1A), numbers on horizontal axes indicate the nucleotide position relative to Wuhan-Hu-1, grey boxes indicate sequence coverage at a certain position, black horizontal lines represent a deletion at a certain position, open reading frames (ORFs) are indicated by colored arrows. A red-lined box indicates the 382-nt deletion. Transcription-regulatory sequences (TRSs) are indicated by yellow arrows.

To investigate the possible effects of the TRS deletion, we calculated and compared the total number of transcripts per million (TPM) of each gene between wild-type (WT) and Δ382 viruses. For each gene, we counted unambiguous reads that uniquely mapped to the gene-specific transcripts that include the joint leader and TRS sequences [9]. We analysed three WT and two Δ382 sequences for which two independent NGS library preparations were available. Our results indicate differential patterns of TPM level between WT and Δ382 viruses (Fig. 2A). The Δ382 sequences displayed a greater level of TPMs in the ORF6, ORF7a and N genes compared to WT. More specifically, the Δ382 N gene, downstream of ORF8, showed a significant increase in the TPM level. To further determine the impact of the 382-nt deletion on the transcription of the N gene, we conducted qPCR to measure the relative abundance of the RNAs from 2 genes (E and N) located upstream and downstream of the deleted region, respectively. Results shown in Figure 2B confirm the enhancement of N gene transcription in Δ382 viruses. This enhancement was consistently observed regardless of the sample type, whether directly from swab samples or passaged viruses.

**Figure 2.**
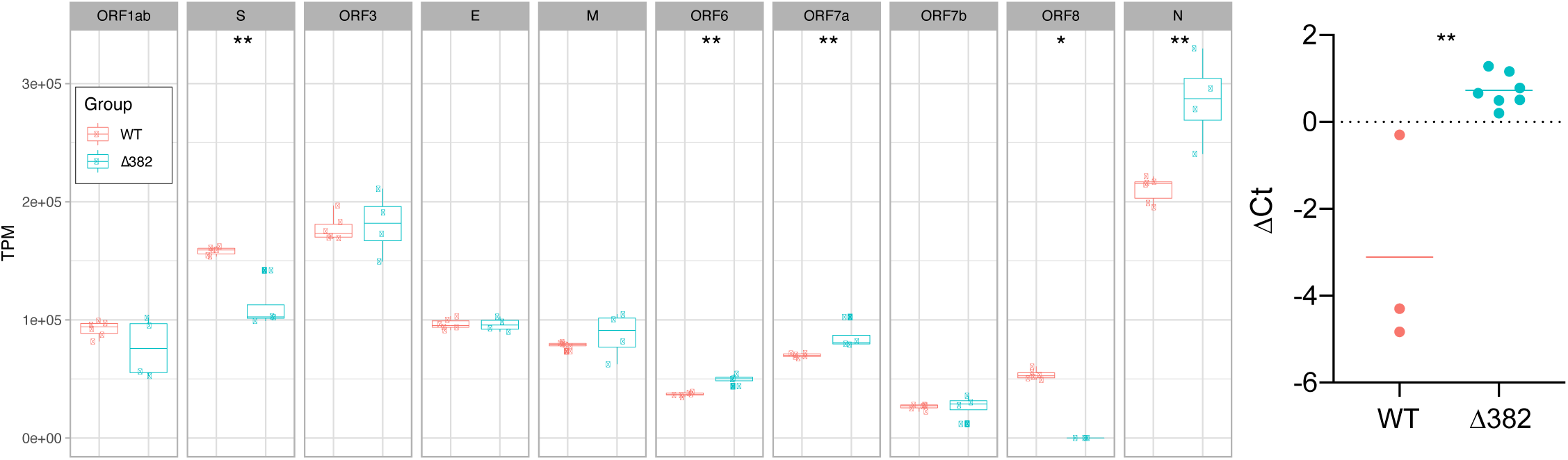
Comparison of transcription of the transcription of the N gene in wild-type (WT) versus Δ382 viruses. **A.** Abundance of mapped reads relative to transcriptional regulatory sequence (TRS) positions across the genome. 70 bp of leader sequence and 230 of each TRS-downstream sequence were merged in a splice-junction in the annotation. Transcripts per million reads (TPM) was calculated from reads mapped specifically to each leader-TRS region and a whisker and scatter plot was drawn for each gene. A Wilcoxon test was applied to the TPM for each gene of Δ382 to WT (*: p ≤ 0.05 **: p ≤ 0.01). **B.** Relative abundance of the RNAs of the E gene in comparison to the N gene of WT versus Δ382 SARS-CoV-2 (p = 0.002) using qPCR. Horizontal bars indicate the mean of each group.

The global phylogeny of SARS-CoV-2 (Suppl. Fig. 2) revealed a comb-like appearance in the phylogenetic tree and a general lack of phylogenetic resolution, reflecting the high similarity of the virus genomes. This tree topology is a typical characteristic of a novel interspecies transmission, wherein the viruses are infecting immunologically naïve populations with sustained human-to-human transmission, as previously shown for the early stages of the 2009 H1N1 pandemic [10]. Notably, we observe an emerging virus lineage (tentatively designated as “Lineage 1” in Suppl. Fig. 2) that consists of sequences from various newly affected countries with recent virus introductions. All viruses from this recent lineage possess an amino acid substitution at residue 250 (from lysine to serine) on the ORF8 gene [11], which is known to undergo strong positive selection during animal-to-human transmission [7].

To estimate the divergence times among lineages of SARS-CoV-2, we reconstructed a dated phylogeny based on 137 complete genomes, including data from this study. The estimated rate of nucleotide substitutions among SARS-CoV-2 viruses is approximately at 8.68×10^−4^ substitutions per site per year (95% HPD: 5.44×10^−4^ –1.22×10^−3^), which is moderately lower than SARS-CoV ([12]: 0.80–2.38×10^−3^) and MERS-CoV (with a mean rate [13] of: 1.12×10^−3^ and 95% HPD: 58.76×10^−4^ –1.37×10^−3^) as well as human seasonal influenza viruses (ranging 1.0–5.5×10^−3^ depending on individual gene segments [10, 14-16]. In contrast, the estimated rate of SARS-CoV-2 viruses is greater than estimates for human coronaviruses by a magnitude order of 4×10^−4^ [17].

The mean TMRCA estimates indicate the introduction of SARS-CoV-2 into humans (Fig. 3. node A) occurred in the middle of November 2019 (95% HPD: 2019.77–2019.94) (Table 1), suggesting that the viruses were present in human hosts approximately one month before the outbreak was detected. A single amino-acid mutation was fixed on the ORF8 region of all full-genome Lineage 1 viruses (Fig. 3, node B), with the TMRCA estimate of 22 December 2019 (95% HPD: 2019.9–2020.0). The Δ382 viruses form a monophyletic group within Lineage 1, and dating estimates indicate that they may have arisen in the human population in Singapore around 07 February 2020 (95% HPD: 2020.12–2020.06), consistent with the date of collection of the Δ382 positive samples (17–19 February 2020). The three Δ382 viruses shared a high level of nucleotide similarities (99.9%) with only 5 nucleotide differences between them. Although the dated phylogeny indicates that Δ382 viruses clustered together and are possibly derived from a single source, there is a general lack of statistical support in SARS-CoV-2 phylogenies due to its recent emergence, and evolutionary inferences should be made with caution.

**Table 1.**
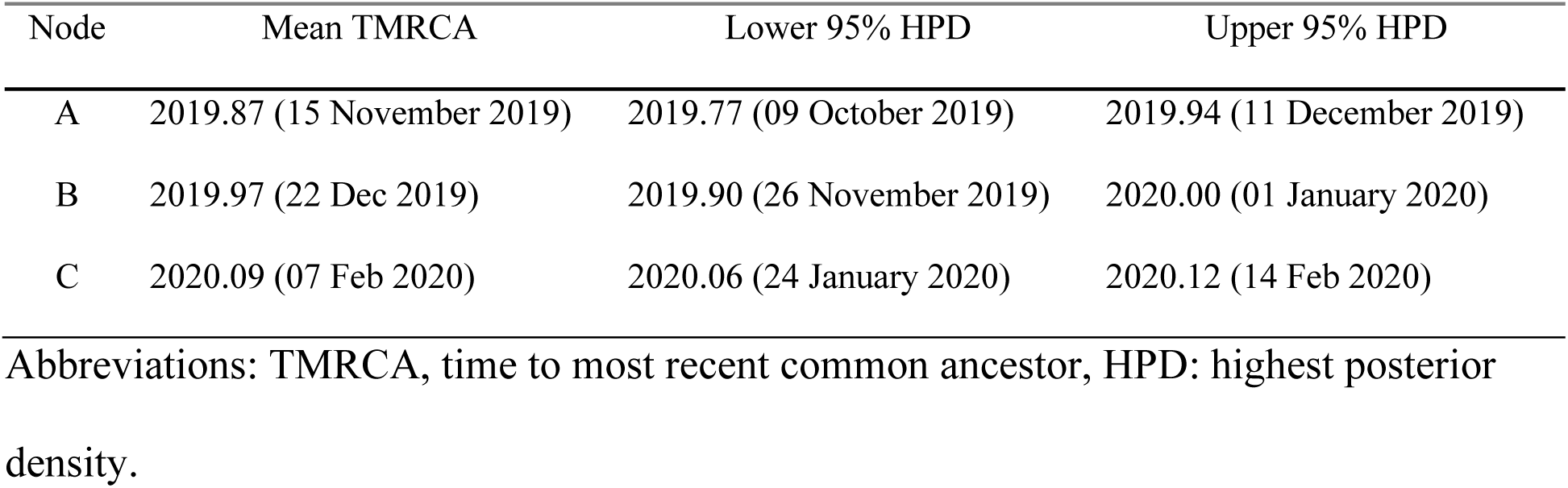
TMRCA estimates of major SARS-CoV-2 nodes.

| Node | Mean TMRCA | Lower 95% HPD | Upper 95% HPD |
| --- | --- | --- | --- |
| A | 2019.87 (15 November 2019) | 2019.77 (09 October 2019) | 2019.94 (11 December 2019) |
| B | 2019.97 (22 Dec 2019) | 2019.90 (26 November 2019) | 2020.00 (01 January 2020) |
| C | 2020.09 (07 Feb 2020) | 2020.06 (24 January 2020) | 2020.12 (14 Feb 2020) |

Abbreviations: TMRCA, time to most recent common ancestor, HPD: highest posterior density.

**Figure 3.**
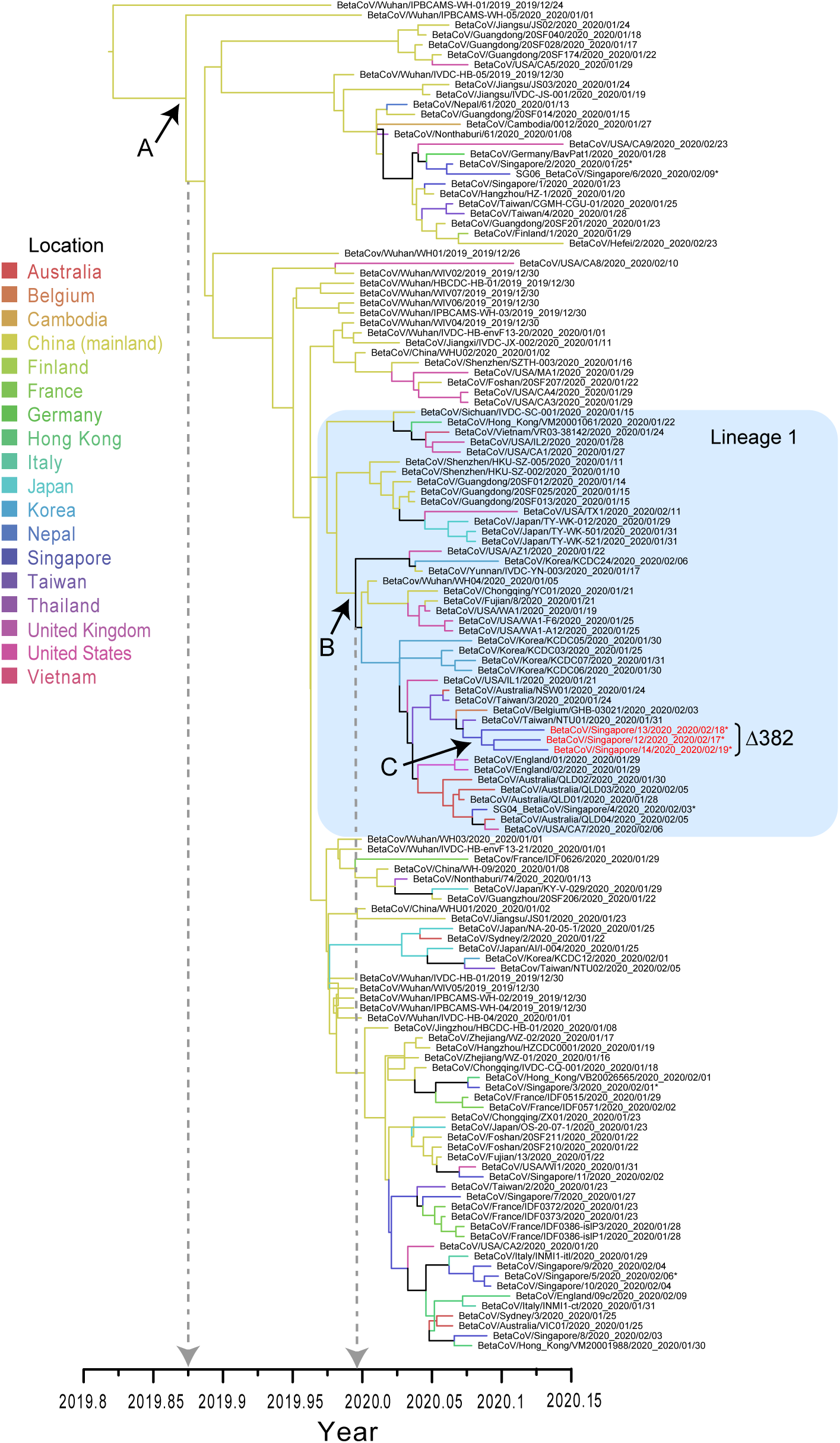
Temporal phylogeny of the complete genomes of SARS-CoV-2 viruses reconstructed using an uncorrelated lognormal relaxed clock model in BEAST. Isolate names in red indicate Δ382 viruses with a large deletion in the ORF7/8 region as described in this study. *Indicates virus genomes generated in this study. Colored branches denote different geographical locations.

In this report we describe the first major evolutionary event of the SARS-CoV-2 virus following its emergence into the human population. Although the biological consequences of this deletion remain unknown, the alteration of the N gene transcription would suggest this may have an impact on virus phenotype. Recent work has indicated that ORF8 of SARS-CoV plays a functional role in virus replicative fitness and may be associated with attenuation during the early stages of human-to-human transmission [2]. Given the prevalence of a variety of deletions in the ORF8 of SARSr-CoVs, it is likely that we will see further deletion variants emerge with the sustained transmission of SARS-CoV-2 in humans. Future research should focus on the phenotypic consequences of Δ382 viruses in the transmission dynamics of the current epidemics and the immediate application of this genomic marker for molecular epidemiological investigation.

## Online content

Any methods, additional references, Nature Research reporting summaries, source data, extended data, supplementary information, acknowledgements, peer review information; details of author contributions and competing interests; and statements of data availability are available at [Article DOI].

## Data availability

The new sequences generated in this study have been deposited in GISAID database under accession numbers EPI_ISL_407987, EPI_ISL_407988, EPI_ISL_410535 to EPI_ISL_410537 for WT viruses and EPI_ISL_414378 to EPI_ISL_414380 for Δ382 viruses.

## Acknowledgements

This study was supported by the Duke-NUS Signature Research Programme funded by the Ministry of Health, Singapore, the National Medical Research Council under its COVID-19 Research Fund (NMRC Project No. COVID19RF-001) and by National Research Foundation Singapore grant NRF2016NRFNSFC002-013 (Combating the Next SARS-or MERS-Like Emerging Infectious Disease Outbreak by Improving Active Surveillance). We thank Viji Vijayan, Benson Ng and Velraj Sivalingam of the Duke-NUS Medical School ABSL3 facility for logistics management and assistance. We thank all scientific staff who assisted with processing clinical samples, especially Velraj Sivalingam, Adrian Kang, Randy Foo, Wan Ni Chia and Akshamal Gamage. We thank Su Ting Tay, Ming Hui Lee and Angie Tan from the Duke-NUS Medical School Genome Biology Facility for expert and rapid technical assistance.

## Author contributions

YCFS, DEA, LFW, GJDS designed and supervised research. BEY, SK, JGHL, LS, GZY, YSL, DCL collected and provided samples. DEA, ML, YZ conducted experiments. YCFS, ZF, JJ, IHM, GJDS performed analyses. YCFS, ZF, ML, LFW, GJDS wrote the paper. All authors reviewed and approved the manuscript.

## Methods

### Ethics statement

This study was undertaken as part of the national disease outbreak and the response and the protocols were approved by the ethics committee of the National Healthcare Group. Patient samples were collected under PROTECT (2012/00917), a multi-centred Prospective Study to Detect Novel Pathogens and Characterize Emerging Infections. Work undertaken at the Duke-NUS Medical School ABSL3 laboratory was approved by the Duke-NUS ABSL3 Biosafety Committee, National University of Singapore and Ministry of Health Singapore.

### Virus culture, RNA extraction and sequencing

Clinical samples from infected patients were collected from public hospitals in Singapore from January–February 2020. Material from clinical samples was used to inoculate Vero-E6 cells (ATCC^®^CRL-1586™). Total RNA was extracted using E.Z.N.A. Total RNA Kit I (Omega Bio-tek) according to manufacturer’s instructions and samples analysed by real-time quantitative reverse transcription-PCR (RT-qPCR) for the detection of SARS-CoV-2 as previously described [18]. Whole genome sequencing was performed using next-generation sequencing (NGS) methodology. The cDNA libraries were constructed using TruSeq RNA Library Prep Kit (Illumina) according to the manufacturer’s instructions and sequenced on an Illumina MiSeq System. Raw NGS reads were trimmed by Trimmomatic v0.39 [19] to remove adaptors and low-quality bases. Genome sequences were assembled and consensus sequences obtained using the BWA algorithm in UGENE v.33. To verify the presence of the deletion in the SARS-CoV-2 genome, we designed two specific PCR primers (F primer: 5’-TGTTAGAGGTACAACAGTACTTT-3’ and R primer: 5’-GGTAGTAGAAATACCATCTTGGA-3’) targeting the ORF7–8 regions. For samples with low Ct values, a hemi-nested PCR was then performed with primers (5’-TGTTTATAACACTTTGCTTCACA-3’) and (5’-GGTAGTAGAAATACCATCTTGGA-3’). The PCR mixture contained the cDNA, primers (10µM each), 10x Pfu reaction buffer (Promega), Pfu DNA polymerase (Promega) and dNTP mix (10mM, Thermo Scientific). The PCR reaction was carried out with the following conditions: 95°C for 2 min, 35 cycles at 95°C for 1 min, 52°C for 30 sec, 72°C for 1 min and a final extension at 72°C for 10 min in a thermal cycler (Applied Biosystems Veriti). Deletions in the PCR products were visualized by gel electrophoresis and confirmed by Sanger sequencing. Three full Δ382 genomes were generated and designated BetaCoV/Singapore/12/2020, BetaCoV/Singapore/13/2020 and BetaCoV/Singapore/14/2020 along with five WT SARS-CoV-2 genomes previously generated in our laboratories and deposited in GISAID (see Suppl. Table 1).

### Genomic characterization

To characterize and map the deletion regions of SARS-CoV-2 viruses, we compared with available SARS-CoV-2 and SARs-CoV related genomes from bat SARs-CoV-JTMC15 (accession number: KY182964) and humans SARs-CoV-Tor 2 (accession number: AY274119) and LC2 (accession number: AY394999). The viral genome organizations of Wuhan-Hu-1 (accession number: MN908947) and Singapore SARS-CoV-2 (Singapore/2 /2020: EPI_ISL_407987) comprise the following gene order and lengths: ORF1ab (open-reading frame) replicase (21291 nt), spike (S: 3822 nt), ORF3 (828 nt), envelope (E: 228 nt), membrane (M: 669 nt), ORF6 (186 nt), ORF7ab (498 nt), ORF8 (366 nt) and nucleocapsid (N: 1260 nt).

To understand possible effects of the TRS deletion on the ORF8 region, we analysed the total number of transcripts per million (TPM) of each gene between wildtype and Δ382 variants. For each gene, unambiguous reads which uniquely mapped to the specific joint leader and TRS sequences are counted. We analysed three patients of SARS-CoV-2 wild type and two patients of SARS-CoV-2 variants and independent NGS library preparations were performed twice for each sample. The Wuhan-Hu-1 was used as the reference sequence for genomic coordinates. Leader RNA sequence is situated at the 5’ UTR region of the genome and the core sequence consists of 26 nt: UCU**CUAAACGAA**CUUUAAAAUCUGUG. Each of the coding genes is preceded by a core transcription regulatory sequence (TRS) which is highly conserved (i.e. ACGAAC), resembling the leader core sequence. The leader RNA is joined to the TRS regions of different genes and the joining is required for driving subgenomic transcription. Coding sequence (CDS), leader and TRS sequences annotations were generated in Geneious and followed published SARS-CoV studies [9]. To characterize the differential levels of TRS for each gene, 70 nt leader sequence and 230 nt downstream of each TRS sequence were annotated individually to form a 300bp leader-TRS transcript for the splicing-aware aligners in R package rtracklayer (v 1.44). NGS raw fastq reads were then mapped to the reference genome by Geneious RNA-Seq mapper with the annotation of splice-junctions for the leader-TRS. For each gene, the TPM of leader-TRS was then calculated in Geneious by excluding ambiguous reads which may come from other TRS. The resulting TPM data was generated in Geneious and graphs plotted using ggplot 2 (v3.2.0) in R v3.6.1. Wilcoxon one-sided tests were performed in R to test the significant differences between WT and Δ382 samples. We also compared individual Ct values of E and N genes between WT and Δ382 viruses using qRT-PCR assays as previously described [18]. Samples used in this analysis included nasopharyngeal swabs from three WT patient samples and both nasopharyngeal swabs and passaged samples from two Δ382 patients (n=7). Significance between groups was assessed using a one-way analysis of variance (ANOVA) in PRISM v8.3.1.

### Phylogenetic analyses

All available genomes of SARS-CoV-2 with associated virus sampling dates were downloaded from the GISAID database. Genome sequence alignment was performed and preliminary maximum likelihood phylogenies of complete genome were reconstructed using RAxML with 1,000 bootstrap replicates in Geneious R9.0.3 software (Biomatters Ltd). Any sequence outliers were removed from subsequent analyses. To reconstruct a time-scaled phylogeny, an uncorrelated lognormal relaxed-clock model with an exponential growth coalescent prior and the HKY85+Γ substitution model was used in the program BEAST v1.10.4 [20] to simultaneously estimate phylogeny, divergence times and rates of nucleotide substitution. Four independent Markov Chain Monte Carlo (MCMC) runs of 100 million generations were performed and sampled every 10,000 generations. The runs were checked for convergence in Tracer v1.7 [21] and that effective sampling size (ESS) values of all parameters was >200. The resulting log and tree files were combined after removing appropriate burn-in values using LogCombiner [20], and the maximum clade credibility (MCC) tree was subsequently generated using TreeAnnontator [20].

---

**Supplementary Figure 1.**
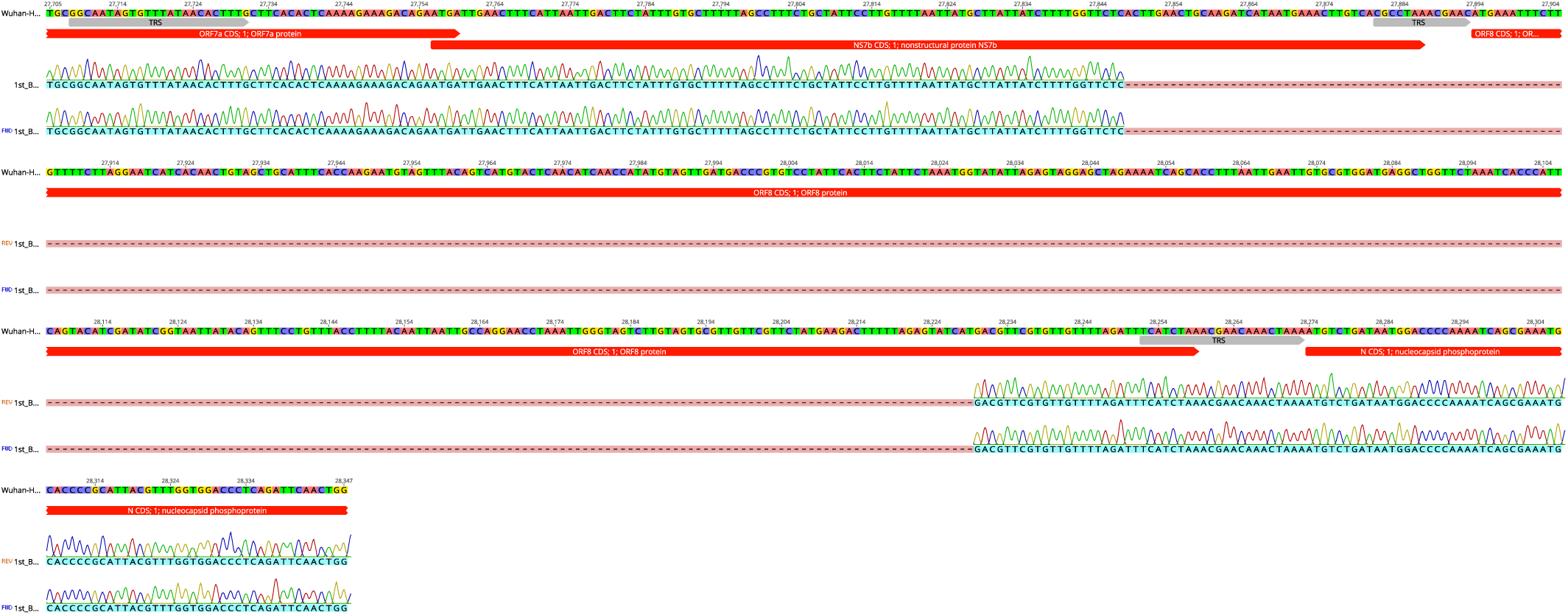
Sanger sequences of BetaCoV/Singapore/12/2020 Δ382 mapped to Wuhan-Hu-1 showing the position of the 382-nt deletion in the SARS-CoV-2 genome.

**Supplementary Figure 2.**
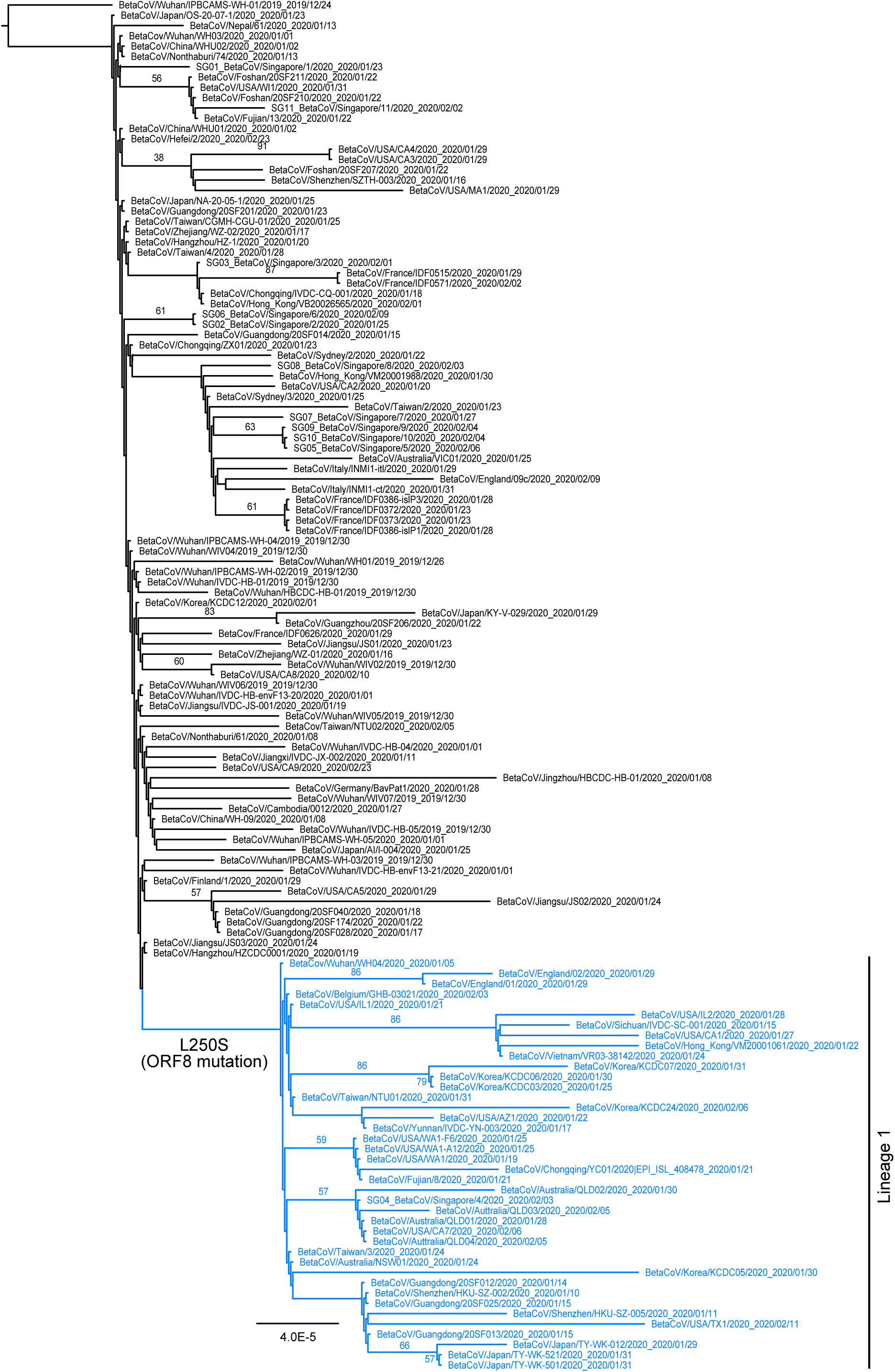
Maximum-likelihood tree of SARS-CoV-2 genomes reconstructed using RAxML with 1,000 bootstrap replicates. Δ382 viruses were exlcluded from this analysis as the L250S mutation is located in the ORF8 deletion.

**Supplementary Table 1.**

| **Accession number** | **Virus isolate** | **Collection date** | **Note** |
| --- | --- | --- | --- |
| EPI_ISL_414378 | BetaCoV/Singapore/12/2020 | 17/02/2020 | 382-nt deletion (this study) |
| EPI_ISL_414379 | BetaCoV/Singapore/13/2020 | 18/02/2020 | 382-nt deletion (this study) |
| EPI_ISL_414380 | BetaCoV/Singapore/14/2020 | 19/02/2020 | 382-nt deletion (this study) |
| EPI_ISL_407987 | BetaCoV/Singapore/2/2020 | 25/01/2020 | Wild type (this study) |
| EPI_ISL_407988 | BetaCoV/Singapore/3/2020 | 01/02/2020 | Wild type (this study) |
| EPI_ISL_410535 | BetaCoV/Singapore/4/2020 | 03/02/2020 | Wild type (this study) |
| EPI_ISL_410536 | BetaCoV/Singapore/5/2020 | 06/02/2020 | Wild type (this study) |
| EPI_ISL_410537 | BetaCoV/Singapore/6/2020 | 09/02/2020 | Wild type (this study) |
| EPI_ISL_402119 | BetaCoV/Wuhan/IVDC-HB-01/2019 | 30/12/2019 | Wild type |
| EPI_ISL_402120 | BetaCoV/Wuhan/IVDC-HB-04/2020 | 01/01/2020 | Wild type |
| EPI_ISL_402121 | BetaCoV/Wuhan/IVDC-HB-05/2019 | 30/12/2019 | Wild type |
| EPI_ISL_402123 | BetaCoV/Wuhan/IPBCAMS-WH-01/2019 | 24/12/2019 | Wild type |
| EPI_ISL_402124 | BetaCoV/Wuhan/WIV04/2019 | 30/12/2019 | Wild type |
| EPI_ISL_402127 | BetaCoV/Wuhan/WIV02/2019 | 30/12/2019 | Wild type |
| EPI_ISL_402128 | BetaCoV/Wuhan/WIV05/2019 | 30/12/2019 | Wild type |
| EPI_ISL_402129 | BetaCoV/Wuhan/WIV06/2019 | 30/12/2019 | Wild type |
| EPI_ISL_402130 | BetaCoV/Wuhan/WIV07/2019 | 30/12/2019 | Wild type |
| EPI_ISL_402132 | BetaCoV/Wuhan/HBCDC-HB-01/2019 | 30/12/2019 | Wild type |
| EPI_ISL_403928 | BetaCoV/Wuhan/IPBCAMS-WH-05/2020 | 01/01/2020 | Wild type |
| EPI_ISL_403929 | BetaCoV/Wuhan/IPBCAMS-WH-04/2019 | 30/12/2019 | Wild type |
| EPI_ISL_403930 | BetaCoV/Wuhan/IPBCAMS-WH-03/2019 | 30/12/2019 | Wild type |
| EPI_ISL_403931 | BetaCoV/Wuhan/IPBCAMS-WH-02/2019 | 30/12/2019 | Wild type |
| EPI_ISL_403932 | BetaCoV/Guangdong/20SF012/2020 | 14/01/2020 | Wild type |
| EPI_ISL_403933 | BetaCoV/Guangdong/20SF013/2020 | 15/01/2020 | Wild type |
| EPI_ISL_403934 | BetaCoV/Guangdong/20SF014/2020 | 15/01/2020 | Wild type |
| EPI_ISL_403935 | BetaCoV/Guangdong/20SF025/2020 | 15/01/2020 | Wild type |
| EPI_ISL_403936 | BetaCoV/Guangdong/20SF028/2020 | 17/01/2020 | Wild type |
| EPI_ISL_403937 | BetaCoV/Guangdong/20SF040/2020 | 18/01/2020 | Wild type |
| EPI_ISL_403962 | BetaCoV/Nonthaburi/61/2020 | 08/01/2020 | Wild type |
| EPI_ISL_403963 | BetaCoV/Nonthaburi/74/2020 | 13/01/2020 | Wild type |
| EPI_ISL_404227 | BetaCoV/Zhejiang/WZ-01/2020 | 16/01/2020 | Wild type |
| EPI_ISL_404228 | BetaCoV/Zhejiang/WZ-02/2020 | 17/01/2020 | Wild type |
| EPI_ISL_404253 | BetaCoV/USA/IL1/2020 | 21/01/2020 | Wild type |
| EPI_ISL_404895 | BetaCoV/USA/WA1/2020 | 19/01/2020 | Wild type |
| EPI_ISL_405839 | BetaCoV/Shenzhen/HKU-SZ-005/2020 | 11/01/2020 | Wild type |
| EPI_ISL_406030 | BetaCoV/Shenzhen/HKU-SZ-002/2020 | 10/01/2020 | Wild type |
| EPI_ISL_406031 | BetaCoV/Taiwan/2/2020 | 23/01/2020 | Wild type |
| EPI_ISL_406034 | BetaCoV/USA/CA1/2020 | 27/01/2020 | Wild type |
| EPI_ISL_406036 | BetaCoV/USA/CA2/2020 | 20/01/2020 | Wild type |
| EPI_ISL_406223 | BetaCoV/USA/AZ1/2020 | 22/01/2020 | Wild type |
| EPI_ISL_406531 | BetaCoV/Guangdong/20SF174/2020 | 22/01/2020 | Wild type |
| EPI_ISL_406533 | BetaCoV/Guangzhou/20SF206/2020 | 22/01/2020 | Wild type |
| EPI_ISL_406534 | BetaCoV/Foshan/20SF207/2020 | 22/01/2020 | Wild type |
| EPI_ISL_406535 | BetaCoV/Foshan/20SF210/2020 | 22/01/2020 | Wild type |
| EPI_ISL_406536 | BetaCoV/Foshan/20SF211/2020 | 22/01/2020 | Wild type |
| EPI_ISL_406538 | BetaCoV/Guangdong/20SF201/2020 | 23/01/2020 | Wild type |
| EPI_ISL_406594 | BetaCoV/Shenzhen/SZTH-003/2020 | 16/01/2020 | Wild type |
| EPI_ISL_406597 | BetaCoV/France/IDF0373/2020 | 23/01/2020 | Wild type |
| EPI_ISL_406716 | BetaCoV/China/WHU01/2020 | 02/01/2020 | Wild type |
| EPI_ISL_406717 | BetaCoV/China/WHU02/2020 | 02/01/2020 | Wild type |
| EPI_ISL_406798 | BetaCov/Wuhan/WH01/2019 | 26/12/2019 | Wild type |
| EPI_ISL_406800 | BetaCov/Wuhan/WH03/2020 | 01/01/2020 | Wild type |
| EPI_ISL_406801 | BetaCov/Wuhan/WH04/2020 | 05/01/2020 | Wild type |
| EPI_ISL_406844 | BetaCoV/Australia/VIC01/2020 | 25/01/2020 | Wild type |
| EPI_ISL_406862 | BetaCoV/Germany/BavPat1/2020 | 28/01/2020 | Wild type |
| EPI_ISL_406970 | BetaCoV/Hangzhou/HZ-1/2020 | 20/01/2020 | Wild type |
| EPI_ISL_406973 | BetaCoV/Singapore/1/2020 | 23/01/2020 | Wild type |
| EPI_ISL_407071 | BetaCoV/England/01/2020 | 29/01/2020 | Wild type |
| EPI_ISL_407073 | BetaCoV/England/02/2020 | 29/01/2020 | Wild type |
| EPI_ISL_407079 | BetaCoV/Finland/1/2020 | 29/01/2020 | Wild type |
| EPI_ISL_407084 | BetaCoV/Japan/AI/I-004/2020 | 25/01/2020 | Wild type |
| EPI_ISL_407193 | BetaCoV/Korea/KCDC03/2020 | 25/01/2020 | Wild type |
| EPI_ISL_407214 | BetaCoV/USA/WA1-A12/2020 | 25/01/2020 | Wild type |
| EPI_ISL_407215 | BetaCoV/USA/WA1-F6/2020 | 25/01/2020 | Wild type |
| EPI_ISL_407313 | BetaCoV/Hangzhou/HZCDC0001/2020 | 19/01/2020 | Wild type |
| EPI_ISL_407893 | BetaCoV/Australia/NSW01/2020 | 24/01/2020 | Wild type |
| EPI_ISL_407894 | BetaCoV/Australia/QLD01/2020 | 28/01/2020 | Wild type |
| EPI_ISL_407896 | BetaCoV/Australia/QLD02/2020 | 30/01/2020 | Wild type |
| EPI_ISL_407976 | BetaCoV/Belgium/GHB-03021/2020 | 03/02/2020 | Wild type |
| EPI_ISL_408008 | BetaCoV/USA/CA3/2020 | 29/01/2020 | Wild type |
| EPI_ISL_408009 | BetaCoV/USA/CA4/2020 | 29/01/2020 | Wild type |
| EPI_ISL_408010 | BetaCoV/USA/CA5/2020 | 29/01/2020 | Wild type |
| EPI_ISL_408430 | BetaCoV/France/IDF0515/2020 | 29/01/2020 | Wild type |
| EPI_ISL_408431 | BetaCov/France/IDF0626/2020 | 29/01/2020 | Wild type |
| EPI_ISL_408478 | BetaCoV/Chongqing/YC01/2020 | 21/01/2020 | Wild type |
| EPI_ISL_408479 | BetaCoV/Chongqing/ZX01/2020 | 23/01/2020 | Wild type |
| EPI_ISL_408480 | BetaCoV/Yunnan/IVDC-YN-003/2020 | 17/01/2020 | Wild type |
| EPI_ISL_408481 | BetaCoV/Chongqing/IVDC-CQ-001/2020 | 18/01/2020 | Wild type |
| EPI_ISL_408484 | BetaCoV/Sichuan/IVDC-SC-001/2020 | 15/01/2020 | Wild type |
| EPI_ISL_408486 | BetaCoV/Jiangxi/IVDC-JX-002/2020 | 11/01/2020 | Wild type |
| EPI_ISL_408488 | BetaCoV/Jiangsu/IVDC-JS-001/2020 | 19/01/2020 | Wild type |
| EPI_ISL_408489 | BetaCoV/Taiwan/NTU01/2020 | 31/01/2020 | Wild type |
| EPI_ISL_408514 | BetaCoV/Wuhan/IVDC-HB-envF13-20/2020 | 01/01/2020 | Wild type |
| EPI_ISL_408515 | BetaCoV/Wuhan/IVDC-HB-envF13-21/2020 | 01/01/2020 | Wild type |
| EPI_ISL_408665 | BetaCoV/Japan/TY-WK-012/2020 | 29/01/2020 | Wild type |
| EPI_ISL_408666 | BetaCoV/Japan/TY-WK-501/2020 | 31/01/2020 | Wild type |
| EPI_ISL_408667 | BetaCoV/Japan/TY-WK-521/2020 | 31/01/2020 | Wild type |
| EPI_ISL_408668 | BetaCoV/Vietnam/VR03-38142/2020 | 24/01/2020 | Wild type |
| EPI_ISL_408669 | BetaCoV/Japan/KY-V-029/2020 | 29/01/2020 | Wild type |
| EPI_ISL_408670 | BetaCoV/USA/WI1/2020 | 31/01/2020 | Wild type |
| EPI_ISL_408976 | BetaCoV/Sydney/2/2020 | 22/01/2020 | Wild type |
| EPI_ISL_408977 | BetaCoV/Sydney/3/2020 | 25/01/2020 | Wild type |
| EPI_ISL_409067 | BetaCoV/USA/MA1/2020 | 29/01/2020 | Wild type |
| EPI_ISL_410045 | BetaCoV/USA/IL2/2020 | 28/01/2020 | Wild type |
| EPI_ISL_410218 | BetaCov/Taiwan/NTU02/2020 | 05/02/2020 | Wild type |
| EPI_ISL_410301 | BetaCoV/Nepal/61/2020 | 13/01/2020 | Wild type |
| EPI_ISL_410531 | BetaCoV/Japan/NA-20-05-1/2020 | 25/01/2020 | Wild type |
| EPI_ISL_410532 | BetaCoV/Japan/OS-20-07-1/2020 | 23/01/2020 | Wild type |
| EPI_ISL_410545 | BetaCoV/Italy/INMI1-itI/2020 | 29/01/2020 | Wild type |
| EPI_ISL_410546 | BetaCoV/Italy/INMI1-ct/2020 | 31/01/2020 | Wild type |
| EPI_ISL_410713 | BetaCoV/Singapore/7/2020 | 27/01/2020 | Wild type |
| EPI_ISL_410714 | BetaCoV/Singapore/8/2020 | 03/02/2020 | Wild type |
| EPI_ISL_410715 | BetaCoV/Singapore/9/2020 | 04/02/2020 | Wild type |
| EPI_ISL_410716 | BetaCoV/Singapore/10/2020 | 04/02/2020 | Wild type |
| EPI_ISL_410717 | BetaCoV/Australia/QLD03/2020 | 05/02/2020 | Wild type |
| EPI_ISL_410718 | BetaCoV/Australia/QLD04/2020 | 05/02/2020 | Wild type |
| EPI_ISL_410719 | BetaCoV/Singapore/11/2020 | 02/02/2020 | Wild type |
| EPI_ISL_410720 | BetaCoV/France/IDF0372/2020 | 23/01/2020 | Wild type |
| EPI_ISL_411060 | BetaCoV/Fujian/8/2020 | 21/01/2020 | Wild type |
| EPI_ISL_411066 | BetaCoV/Fujian/13/2020 | 22/01/2020 | Wild type |
| EPI_ISL_411218 | BetaCoV/France/IDF0571/2020 | 02/02/2020 | Wild type |
| EPI_ISL_411219 | BetaCoV/France/IDF0386-isIP1/2020 | 28/01/2020 | Wild type |
| EPI_ISL_411220 | BetaCoV/France/IDF0386-isIP3/2020 | 28/01/2020 | Wild type |
| EPI_ISL_411902 | BetaCoV/Cambodia/0012/2020 | 27/01/2020 | Wild type |
| EPI_ISL_411915 | BetaCoV/Taiwan/CGMH-CGU-01/2020 | 25/01/2020 | Wild type |
| EPI_ISL_411926 | BetaCoV/Taiwan/3/2020 | 24/01/2020 | Wild type |
| EPI_ISL_411927 | BetaCoV/Taiwan/4/2020 | 28/01/2020 | Wild type |
| EPI_ISL_411950 | BetaCoV/Jiangsu/JS01/2020 | 23/01/2020 | Wild type |
| EPI_ISL_411952 | BetaCoV/Jiangsu/JS02/2020 | 24/01/2020 | Wild type |
| EPI_ISL_411953 | BetaCoV/Jiangsu/JS03/2020 | 24/01/2020 | Wild type |
| EPI_ISL_411954 | BetaCoV/USA/CA7/2020 | 06/02/2020 | Wild type |
| EPI_ISL_411955 | BetaCoV/USA/CA8/2020 | 10/02/2020 | Wild type |
| EPI_ISL_411956 | BetaCoV/USA/TX1/2020 | 11/02/2020 | Wild type |
| EPI_ISL_411957 | BetaCoV/China/WH-09/2020 | 08/01/2020 | Wild type |
| EPI_ISL_412026 | BetaCoV/Hefei/2/2020 | 23/02/2020 | Wild type |
| EPI_ISL_412028 | BetaCoV/Hong_Kong/VM20001061/2020 | 22/01/2020 | Wild type |
| EPI_ISL_412029 | BetaCoV/Hong_Kong/VM20001988/2020 | 30/01/2020 | Wild type |
| EPI_ISL_412030 | BetaCoV/Hong_Kong/VB20026565/2020 | 01/02/2020 | Wild type |
| EPI_ISL_412116 | BetaCoV/England/09c/2020 | 09/02/2020 | Wild type |
| EPI_ISL_412459 | BetaCoV/Jingzhou/HBCDC-HB-01/2020 | 08/01/2020 | Wild type |
| EPI_ISL_412862 | BetaCoV/USA/CA9/2020 | 23/02/2020 | Wild type |
| EPI_ISL_412869 | BetaCoV/Korea/KCDC05/2020 | 30/01/2020 | Wild type |
| EPI_ISL_412870 | BetaCoV/Korea/KCDC06/2020 | 30/01/2020 | Wild type |
| EPI_ISL_412871 | BetaCoV/Korea/KCDC07/2020 | 31/01/2020 | Wild type |
| EPI_ISL_412872 | BetaCoV/Korea/KCDC12/2020 | 01/02/2020 | Wild type |
| EPI_ISL_412873 | BetaCoV/Korea/KCDC24/2020 | 06/02/2020 | Wild type |

## References

1. Chinese, S.M.E.C., Molecular evolution of the SARS coronavirus during the course of the SARS epidemic in China. Science, 2004. 303(5664): p. 1666–9.

2. Muth, D., et al., Attenuation of replication by a 29 nucleotide deletion in SARS-coronavirus acquired during the early stages of human-to-human transmission. Sci Rep, 2018. 8(1): p. 15177.

3. Lu, R., et al., Genomic characterisation and epidemiology of 2019 novel coronavirus: implications for virus origins and receptor binding. The Lancet, 2020. 395(10224): p. 565–574.

4. World_Health_Organization. Coronavirus disease (COVID-2019) situation reports. 2020 [cited 2020 Mar 2]; Available from: https://www.who.int/emergencies/diseases/novel-coronavirus-2019/situation-reports.

5. Chinese, S.M.E.C., Molecular evolution of the SARS coronavirus during the course of the SARS epidemic in China. Science (New York, N.Y.), 2004. 303(5664): p. 1666–1669.

6. Wang, L.-F., et al., Review of bats and SARS. Emerging infectious diseases, 2006. 12(12): p. 1834–1840.

7. Lau, S.K., et al., Severe Acute Respiratory Syndrome (SARS) Coronavirus ORF8 Protein Is Acquired from SARS-Related Coronavirus from Greater Horseshoe Bats through Recombination. J Virol, 2015. 89(20): p. 10532–47.

8. Xu, L., et al., Detection and characterization of diverse alpha- and betacoronaviruses from bats in China. Virol Sin, 2016. 31(1): p. 69–77.

9. Hussain, S., et al., Identification of novel subgenomic RNAs and noncanonical transcription initiation signals of severe acute respiratory syndrome coronavirus. J Virol, 2005. 79(9): p. 5288–95.

10. Su, Y.C.F., et al., Phylodynamics of H1N1/2009 influenza reveals the transition from host adaptation to immune-driven selection. Nature Communications, 2015. 6: p. 7952.

11. Tang, X., et al., On the origin and continuing evolution of SARS-CoV-2. National Science Review, 2020.

12. Zhao, Z., et al., Moderate mutation rate in the SARS coronavirus genome and its implications. BMC Evolutionary Biology, 2004. 4(1): p. 21.

13. Cotten, M., et al., *Spread, circulation*, and evolution of the Middle East respiratory syndrome coronavirus. mBio, 2014. 5(1): p. e01062–13.

14. Vijaykrishna, D., et al., The contrasting phylodynamics of human influenza B viruses. eLife, 2015. 4: p. e05055.

15. Virk, R.K., et al., Divergent evolutionary trajectories of influenza B viruses underlie their contemporaneous epidemic activity. Proceedings of the National Academy of Sciences, 2020. 117(1): p. 619–628.

16. Rambaut, A., et al., The genomic and epidemiological dynamics of human influenza A virus. Nature, 2008. 453(7195): p. 615–619.

17. Vijgen, L., et al., Complete Genomic Sequence of Human Coronavirus OC43: Molecular Clock Analysis Suggests a Relatively Recent Zoonotic Coronavirus Transmission Event. Journal of Virology, 2005. 79(3): p. 1595–1604.

18. Corman, V.M., et al., Detection of 2019 novel coronavirus (2019-nCoV) by real-time RT-PCR. Eurosurveillance, 2020. 25(3): p. 2000045.

19. Bolger, A.M., M. Lohse, and B. Usadel, Trimmomatic: a flexible trimmer for Illumina sequence data. Bioinformatics, 2014. 30(15): p. 2114–2120.

20. Suchard, M.A., et al., Bayesian phylogenetic and phylodynamic data integration using BEAST 1.10. Virus Evol, 2018. 4(1): p. vey016.

21. Rambaut, A., et al., Posterior Summarization in Bayesian Phylogenetics Using Tracer 1.7. Syst Biol, 2018. 67(5): p. 901–904.

